# A standard operating procedure for outlier removal in large-sample epidemiological transcriptomics datasets

**DOI:** 10.1101/144519

**Authors:** Hege Marie Bøvelstad, Einar Holsbø, Lars Ailo Bongo, Eiliv Lund

## Abstract

Transcriptome measurements and other –omics type data are increasingly more used in epidemiological studies. Most of omics studies to date are small with samples sizes in the tens, or sometimes low hundreds, but this is changing. Our Norwegian Woman and Cancer (NOWAC) datasets are to date one or two orders of magnitude larger. The NOWAC biobank contains about 50000 blood samples from a prospective study. Around 125 breast cancer cases occur in this cohort each year. The large biological variation in gene expression means that many observations are needed to draw scientific conclusions. This is true for both microarray and RNA-seq type data. Hence, larger datasets are likely to become more common soon.

Technical outliers are observations that somehow were distorted at the lab or during sampling. If not removed these observations add bias and variance in later statistical analyses, and may skew the results. Hence, quality assessment and data cleaning are important. We find common quality assessment libraries difficult to work with for large datasets for two reasons: slow execution speed and unsuitable visualizations.

In this paper, we present our standard operating procedure (SOP) for large-sample transcriptomics datasets. Our SOP combines automatic outlier detection with manual evaluation to avoid removing valuable observations. We use laboratory quality measures and statistical measures of deviation to aid the analyst. These are available in the *nowaclean* R package, currently available on GitHub (https://github.com/3inar/nowaclean). Finally, we evaluate our SOP on one of our larger datasets with 832 observations.

## 1 Introduction

The use of –omics data in epidemiological studies is now common. Typical studies comprise sample sizes in the tens or low hundreds, but sizes in the order of thousands will soon be common. The NOWAC postgenome cohort (1) contains blood samples from 50000 women. In this cohort there are approximately 125 new breast cancer cases per year, and we have thus far extracted and processed blood samples from 1660 case—control pairs, or 3320 blood samples in total. Omics experiments are elaborate procedures with several steps. In the case of microarrays, these include mRNA isolation, hybridization, washing, and scanning. Each step may add random or systematic errors. Technical errors may also come from supplies or instruments. Mishaps may occur in the lab. The samples themselves can get contaminated in various ways. In whole-blood samples, there is also the added challenge of mRNA degradation due to high RNase activity. All this may be detrimental to the quality of the data and hence affect downstream analyses.

The goal of gene expression experiments is to detect differences in gene expression levels between groups. This is usually evaluated gene-by-gene or for sets of related genes. The methods for such analyses depend on accurate estimation of the sample variance. If there are technical outliers contributing unnecessary variance, removing these should increase power. However, removing biological outliers will result in underestimation of the natural biological variance. This in turn will increase the risk of spurious conclusions. There is a fine line to tread, and the accurate identification of technical outliers is important for later analysis.

Many publications guide the identification of outliers in gene expression data (2–5). Yet, there is no real consensus on the best approach. For example, some authors such as (6) propose automated procedures for outlier removal. Others such as (4) warn against automation and instead recommend careful investigation.

Outlier removal is particularly challenging for studies based on blood samples, since there is larger biological variation in gene expression data from blood than in tumor tissue (7). The strength of the signal in tumor tissue makes it much more robust to variance than the signal in blood samples, which is weak and variable. This makes it more difficult to distinguish outliers from non-outliers and signal from noise. It is not well known whether lifestyle factors like medication use affect blood gene expression. All this complicates outlier identification, and it’s inadvisable to remove outliers in a systematic, automated way.

R-packages such as *arrayQualityMetrics* (AQM) (3) and *lumi* (8) implement the most popular outlier detection methods for gene expression data. Important to these approaches is the combination of computational methods with interactive visualization. However, when dealing with several hundreds of observations, these methods are cumbersome for two reasons. First, some methods are slow and thus inefficient for interactive use. Second, their visualizations do not work well for larger sample sizes due to overplotting. The latter is in our opinion the most important aspect, as the decision to remove an outlier often rests on visual inspection by the analyst.

We also wish to provide numerical measures and standardized guidelines to help the user. The measures are statistics of deviation, derived from the data, and laboratory quality metrics. We believe that standardization removes some of the subjectivity from the task. Standardizing the outlier removal procedure as much as possible will enhance reproducibility and consistency.

Below we describe our standard operating procedure (SOP) for outlier removal in large-sample transcriptomics datasets. We believe our SOP will strengthen the reporting of observational studies in epidemiology (9). We have implemented the SOP as an open source R package that combines automated outlier detection with expert evaluation. The automated part consists of ranking observations by deviation metrics. We base these metrics on standard methods for outlier removal in data from microarrays. Our improvements are faster execution and easier-to-read visualizations.

We provide a unified, interactive interface, saving computations for tinkering with thresholds and using standard R methods where available. We use data from the NOWAC study (10) for demonstration and evaluation.

Our R package is open source and available at:

https://github.com/3inar/nowaclean

## 2 Methods

### 2.1 Data

The NOWAC study is a nation-wide, population-based cancer study (1). A thorough description of the NOWAC postgenome cohort can be found in (10). To summarize: 97.2% of the women in the NOWAC cohort consented to donate a blood sample to research. Out of these, about two thirds ended up providing an actual blood sample. Blood sampling kits were sent out in batches of 500. These kits included a two-page questionnaire and a PAXgene tube (PreAnalytiX GmbH, Hembrechtikon, Switzerland). For the most part, the family general practitioner drew the actual blood sample. The sample was then mailed overnight to Tromsø. Between 2003 and 2006 the NOWAC biobank grew to comprise 48,692 blood samples. These make up the NOWAC postgenome cohort. The Norwegian Cancer Registry provides yearly updates about cancer cases. Statistics Norway provides yearly updates about emigrations and deaths. A control sample is assigned to each breast cancer case in the cohort yielding a nested case-control design. These are matched on mailing batch, time of blood sampling and year of birth. We keep each case—control pair together through every step in the laboratory. The statistical analysis of microarray data is described in (11).

In this paper, we use a subset of 832 observations from the NOWAC cohort. The Genomics Core Facility at the Norwegian University of Science and Technology provided the laboratory work. They processed the samples on Illumina Whole-Genome Gene Expression Bead Chips (http://technology.illumina.com/technology/beadarray-technology.html), HumanHT-12 v4. The raw microarray images are processed in GenomeStudio (http://bioinformatics.illumina.com/informatics/sequencing-microarray-data-analysis/genomestudio.html). This is Illumina’s own software for processing data from their platforms. The result is a table of 47323 probes for 832 observations on the summary level: one number per probe per observation.

### 2.2 The NOWAC pipeline

The outlier SOP is part of our data processing pipeline in NOWAC. The pipeline (Fig. 1) contains three major steps where outlier removal is Step 2. We briefly describe the data preparation (Step 1) and the preprocessing (Step 3) to provide context for the SOP.

**Fig. 1.**
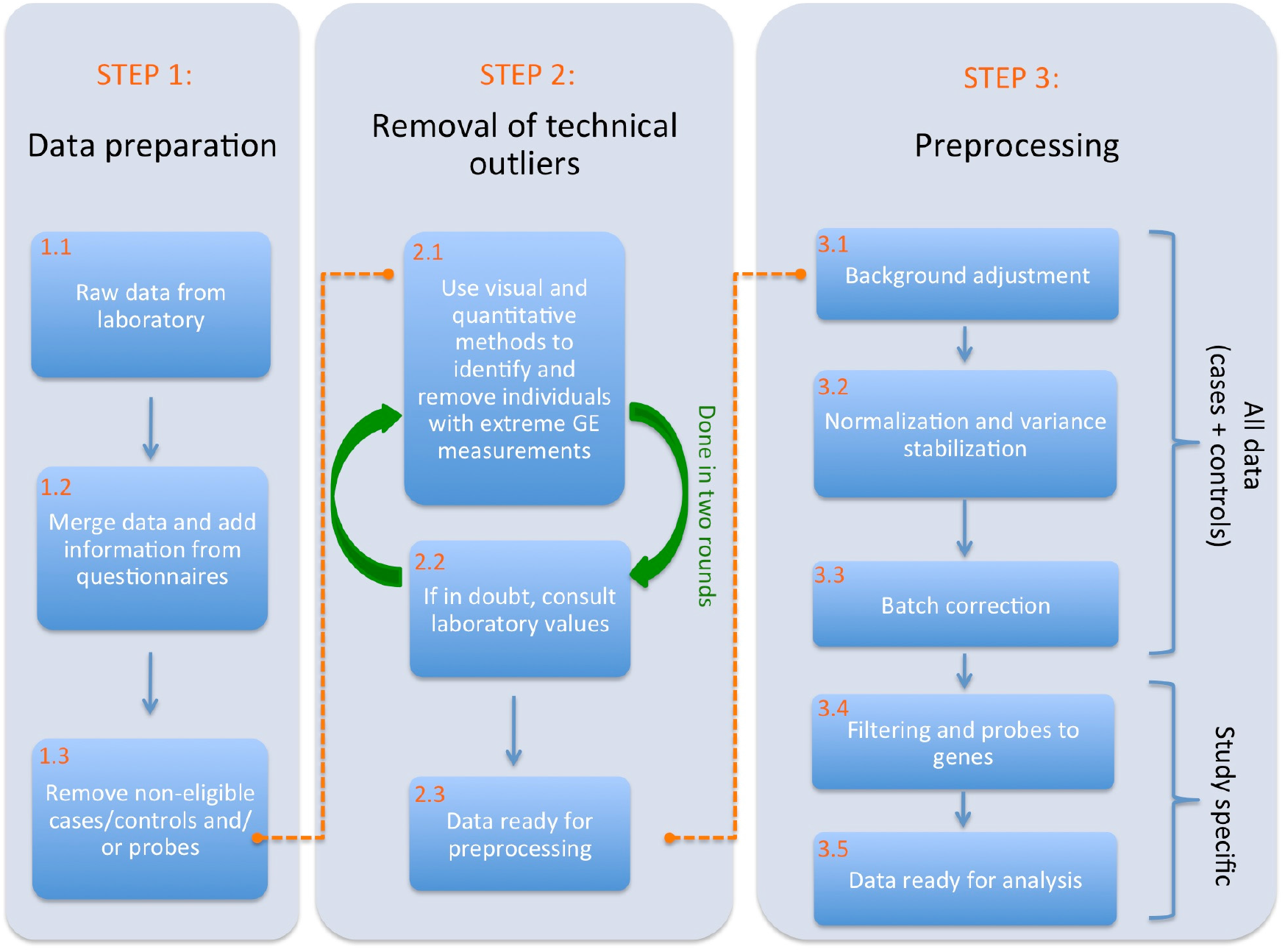
*NOWAC (10) standardized data analysis pipeline for cleaning and preprocessing the data. The pipeline is split into three steps, where all steps up to and including Step 3.3 are performed using all cases and controls that are eligible and not considered as outliers. Step 3.4 needs to be performed for each specific study/question at hand, fine-tuning the data to optimize the power of the analysis. Abbreviation: GE=Gene expression.*

**STEP 1.1:** Described in the “Data” section above.

**STEP 1.2:** Microarray gene expression measurements from the lab are merged into an R LumiBatch-object (8) based on a unique lab number, along with external information from questionnaires, the Norwegian Cancer Registry, and Statistics Norway.

**STEP 1.3:** Yearly updates from the Cancer Registry can reveal that controls have become cases, or that cases have received a second cancer diagnosis. We considered these individuals non-eligible, and remove them along with their matching case/control.

For multivariate analyses, we remove 38 probes related to blood type, specifically the human leukocyte antigen (HLA) system. These are usually expressed strongly and have high variance, which will affect multivariate analyses. We have seen that they can dominate the variance-covariance pattern in the principal component analysis (PCA) transformation of the data (will be described in detail in the next section), and as such other patterns might be obscured. This is relevant for our SOP as we do PCA, so we recommend to take these out before outlier detection. It is possible to put these probes back after outlier detection. The decision will depend on whether the genes are interesting for subsequent analyses.

**STEP 2:** Described in detail below in the “Outlier Removal SOP” section.

**STEPS 3.1 and 3.2:** We apply the normal-exponential background adjustment method to make signals comparable across individuals (12,13). We also use quantile normalization (14) and log2-transform the data to stabilize the variance.

**STEP 3.3:** Batch effects are systematic errors introduced when processing blood samples in multiple batches in the laboratory. Examples of a batch are all chips that are processed at the same day (named plate), laboratory technician, or the batch of laboratory regents used. It’s important to adjust for batch effects with methods like e.g. ComBat (15).

**STEP 3.4:** We filter out probes that are likely to be below the level of detection, probes that are expressed in only a few arrays, and probes that are known to have unreliable annotation. This reduces the number of probes and the risk of false positives in subsequent analysis.

### 2.3 Outlier removal SOP

Outlier removal is a subjective task. Our SOP combines guidelines, visualizations, and quantitative measures to help. An outlier should only be excluded if it is of technical origin, since biological outliers are valuable. Technical measures from the microarray lab describe the quality of the blood sample. They include information on RNA abundance, mRNA contamination, etc. They may provide hints to why an array might look wrong and help make the distinction between technical outliers and biological outliers. We describe the lab measures in detail further below. As extreme outliers have a strong influence on many of the plots and measures we use, we do outlier detection/removal in two rounds. In case—control designs, when an outlying observation is removed, the matching case/control will also be removed. An overview of the SOP is provided in **Fig. 2.**

**Fig. 2:**
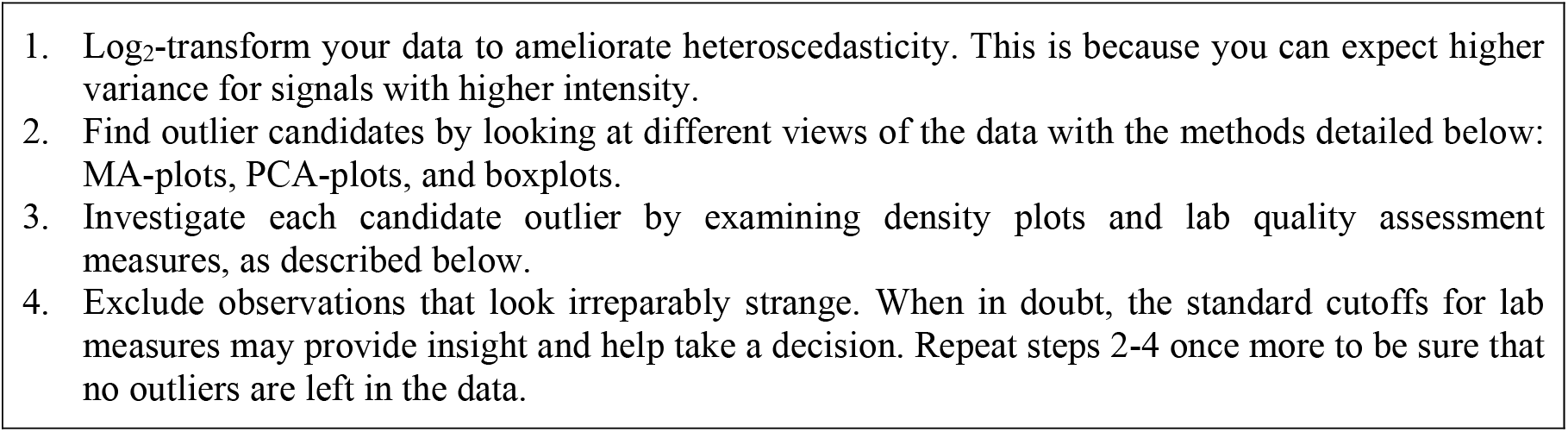
*Our outlier removal standard operating procedure*

We evaluate **individual array quality** with array-wise MA-plots (16) where we compare each array with the median array. An MA-plot is a mean—difference plot that compares two assays on the log2 scale. Specifically, let A_1_ be a given array, and A_2_ the median array constructed by taking gene-wise medians over all arrays. Then, compute the two statistics: 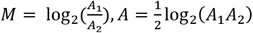.

You should expect M to be constant as a function of A in well-behaved arrays (**Fig. 3**, bottom panel). A trend in M as a function of A would indicate that gene expression values are somehow systematically skewed away from the median array (**Fig. 3**, top panel).

**Fig. 3.**
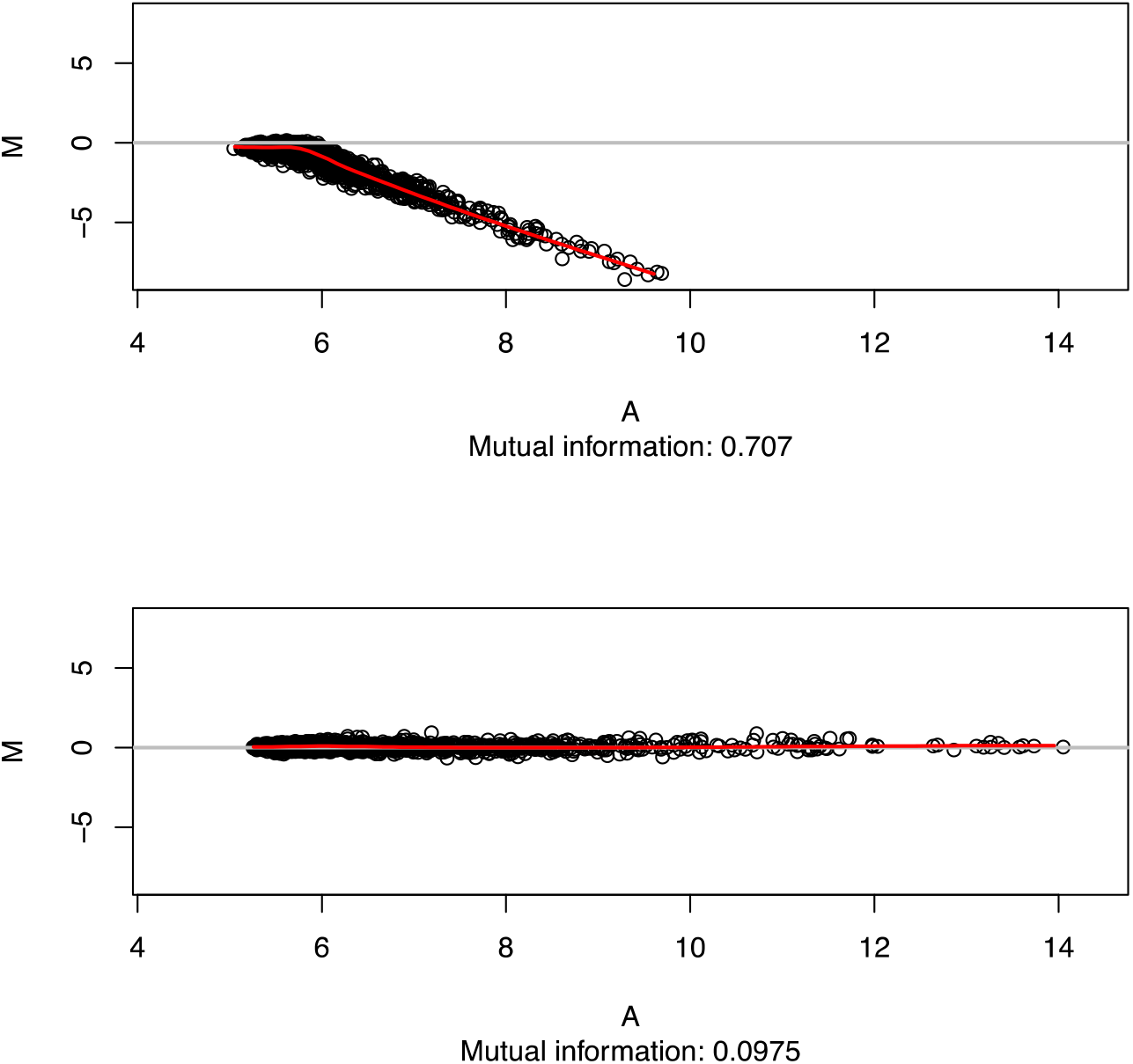
*Illustrations of MA plots. On top: a potential outlier array. On bottom: a well-behaved array.*

You can measure independence of M and A in several ways, AQM uses Hoeffding’s D statistic, which measures squared deviance from independence (difference between joint density and product of marginal densities). We use the similar measure of mutual information (MI), defined as

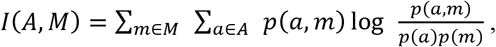

simply because the R-implementation is considerably faster (17). The joint density of the two statistics must be discretized, so some information may be lost. For this and many other reasons it’s important to inspect the outliers yourself.

We evaluate **homogeneity between arrays** by inspecting boxplots. As we have hundreds to thousands of arrays, doing regular boxplots will result in overplotting. Hence, we use “compressed” boxplots where each quantile is represented by a single point, and the points for corresponding quantiles are connected by lines. The same is implemented for the lower and upper whiskers of the boxplot, given by the most extreme data point within 1.5 times the interquartile range from the median. This results in a plot with five continuous horizontal lines (**Fig. 4**). We measure deviation from normal data by comparing the empirical cumulative distribution function (ECDF) of the expression intensities for each array with the ECDF of all arrays pooled. Distance from the pooled ECDF is measured by the Kolmogorov-Smirnov (KS) statistic (18), which measures the largest distance between two distribution functions. By default, we order the boxplots by their respective KS statistic, but it may also be interesting to order by other things such as plate number to look for batch effects.

**Fig. 4.**
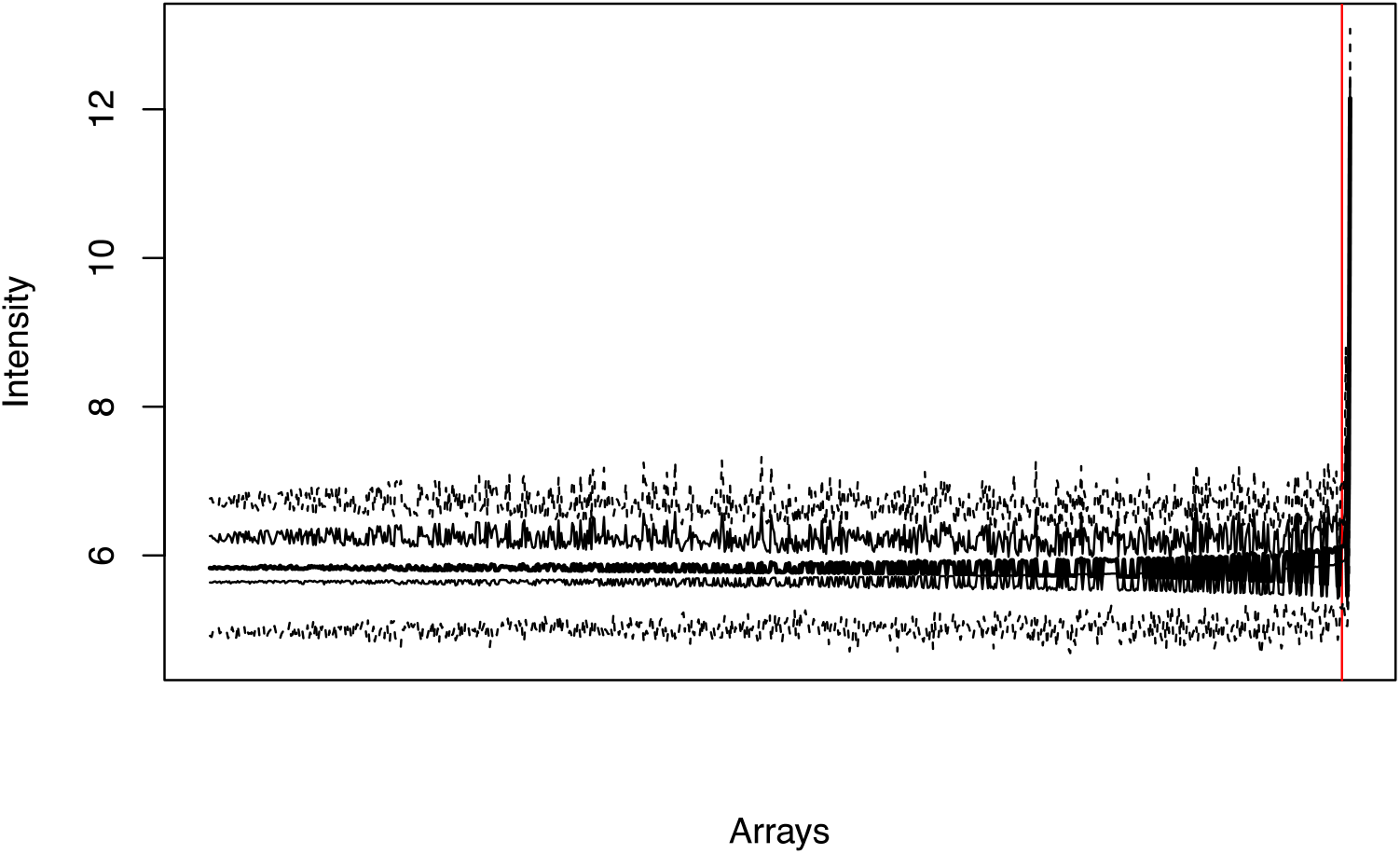
*Boxplots, ordered by the KS-statistics. The individuals with a boxplot to the right of the red line has a KS-statistics above m=3 standard deviations from the mean observed KS-statistics. Points in this plot represent the components of a boxplot: median, lower and upper quartiles, lower and upper whiskers. Points of the same type (e.g. median) are joined by a line.*

We define outliers as those observations that fall outside *m* standard deviations from the mean observed KS statistic. The value of *m* will depend on how conservative the analyst is in its search for outliers. The higher value of *m*, the fewer individuals will be marked as outliers. For the example shown in **Fig. 4** we used *m = 3 (indicated by a red line).*

For **between-array comparison** we apply principal component analysis (PCA) (19) to the data, and display the first two principal components in a scatterplot. As a quantitative measure to guide the outlier identification, we compute the Mahalanobis distance of all arrays to the mean array. In PCA-transformed data the Mahalanobis distance and Euclidean distance are necessarily equal. As shown in the top panel of **Fig. 5**. there can be a distinct shape and rotation to the data that’s not captured due to outliers. Hence, we define a “central cluster” that’s used to compute distances. We obtain this central cluster by ignoring points that are less likely than 99% (adjustable) by Chebyshev’s inequality. This leads to distances that fit the shape and rotation of the data better, see the bottom panel of **Fig. 5**. Outliers are then defined as those more than *n* standard deviations from the data center in Mahalanobis distance. Once again, the value of *n* must be determined by the analyst; We used *n = 3* in the analysis below.

**Fig. 5.**
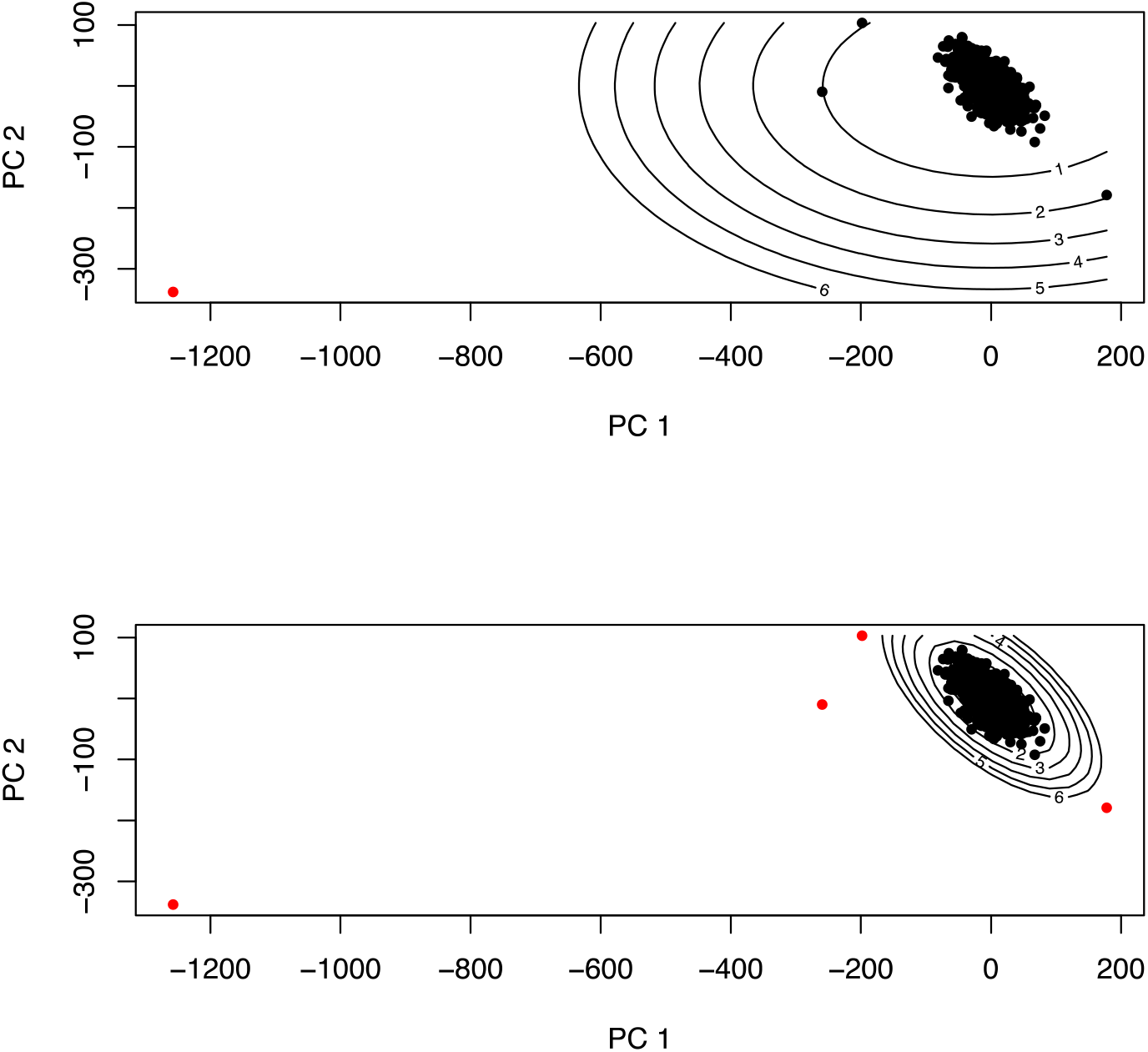
*PCA plots. The lines show Mahalanobis distance to the center of the data (in standard deviations). The red points are considered potential outliers as they are farther away than two standard deviations. The top panel shows the distances computed from all data points. The bottom panel shows the same when leaving out the least likely points by Chebyshev’s inequality.*

Finally, we use density plots to inspect observations we suspect to be technical outliers based on the methods described above. These plots show distribution properties that are hidden in the other plots like severely skewed modes or several modes, neither of which you should expect to see in well-behaved data. **Fig. 7** below in the Results section shows an example of a density plot.

After exploring different outlier detection methods, the analyst is left with a selection of outliers and must decide which are technical outliers that should be discarded. Several **technical measures from the laboratory may help** guide this decision. These measures include information on RNA abundance; the quality of the blood sample in terms of mRNA degradation, quantified by RNA Integrity Number (RIN); and the level of contamination in the blood sample, quantified by NanoDrop 260/230 and 280/230 ratios. These values may help the analyst understand why some observations have outlying values. For borderline outliers, where the analyst is uncertain, we provide standard exclusion thresholds for each lab measure (**Fig. 6**). If the observation is suspect and it falls outside of any of these thresholds, it may be regarded as an outlier and thus discarded. It is entirely possible for observations to look perfectly sensible despite bad lab measures, hence we don’t exclude observations based purely on these numbers. We consult the lab measures only once we suspect an array to be a technical outlier based on the plots.

**Fig. 6.**
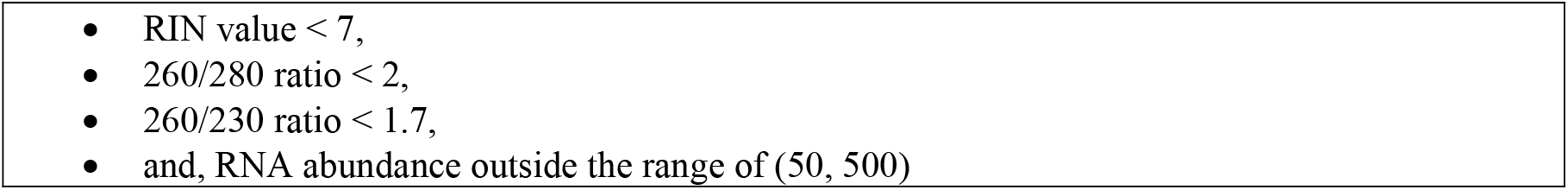
Standard exclusion thresholds for technical measures from the laboratory

### 2.4 The *nowaclean* R Package

The nowaclean R package (for details and source code, see https://github.com/3inar/nowaclean) implements our standard operating procedure for detecting and removing technical outliers in the NOWAC microarray data. As mentioned above, the functionality we provide already exists elsewhere. The novelty of this R-package is the improved speed and visualizations for data sets with a large number of arrays.

We have through this work identified four design principles that we believe improve the user experience: i) save computations so that users can tune thresholds; ii) force the use of names instead of indices into a matrix, in case several representations of the data are in use; iii) have a unified interface to the different methods: always use R’s standard *predict* and *plot* methods, and provide the same set of arguments to these as far as possible; and finally iv) decouple the methods from special types of objects such as the Bioconductor standard *esets* and work on built-in matrices instead. This last point is to provide functionality to a broader user base.

### 2.5 Evaluation methods for outlier removal

To study how our SOP affects downstream analysis we need to quantify the effect of the outlier removal. As removing individuals may reduce power, removing technical outliers identified in our SOP should on the contrary increase the power and make sure that the downstream analysis leads to more sound and biologically reliable results. One way to quantify the effect of the SOP outlier removal is to count the number of genes that are differentially expressed between cases and controls before and after outlier removal, as described in (6). A gene *G* is differentially expressed between cases and controls if *μ(G_cases_) ≠ μ(G_controls_)*, i.e. the average expression *μ(G)* of gene *G* in one group is different from that of the other, as determined by the *limma* moderated t-test (20).

We will examine the number of significant findings in two situations. First in a situation where we know there is no difference between groups. We create a pseudosample by assigning observations to groups randomly to ensure no relationship between group and gene expression levels (i.e. permutation). In this pseudosample we compute the statistic 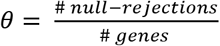, i.e. the proportion of significant genes. To get a distribution over *θ*, we will repeat the procedure for 1500 random pseudosamples. In this situation there is no difference between groups, and thus the null hypothesis should be rejected for about 5% of all genes tested when using a significance level of *α = 0.05.* Hence *θ* is in effect an estimate of type-I error rate, the effective size of the hypothesis test.

In the second situation, we count the number of rejected null hypotheses when we expect a difference between groups. We will use an anonymized group variable from the NOWAC questionnaires, and generate data where we draw (observation, group) pairs with replacement from the real data, replicating the original dataset size (i.e. bootstrapping). We then compute the statistic *θ* defined above, and repeat the procedure for 1000 bootstrap samples. By doing this we get an estimate of the variance of *θ*, and not just a point estimate. In this situation *θ* says something about statistical power, but is not a direct estimate. We are primarily interested in a comparison of outlier removal strategies, so change in *θ* is what is most important.

The fraction of outliers to non-outliers is likely to be small, and their effect might be subtle in large samples. For this reason, we will inspect three dataset sizes: all of our data, half the data, and 10% of the data. We do this by, for each new pseudosample, removing the correct fraction of observations from the full data set but making sure that the identified outliers are kept in the pseudosample. This is done for both the permutation and the bootstrapping experiment. We then perform the preprocessing described in STEP 3 of **Fig. 1**, and finally compute *θ* for the data with and without outliers.

As for any procedure involving hypothesis testing you ideally want as high a statistical power as possible, and it’s nice if you get the correct test size. That is, you want as many type-I errors as you’d expect so that *α = θ* at the *α* level.

## 3 Results

We demonstrate our methods on a typical NOWAC raw data set comprising 47323 probes for 832 observations. After applying our SOP, we have identified four observations as technical outliers. We describe this process in detail in **Online Resource 1**. We used version 0.2.8 of *nowaclean* for these computations. The source code is available as **Online Resource 2**, although it is simpler to install from the instructions at https://github.com/3inar/nowaclean.

**Fig. 7** shows the expression densities of the four outliers (red lines) along with all the other observations (black lines). If by some chance the three right-skewed observations are e.g. all cases and the left-skewed observation is a control, the result would almost certainly be overestimation of differential expression. These same four observations are the ones highlighted in red above in the PCA plot of **Fig. 5**.

**Fig. 7:**
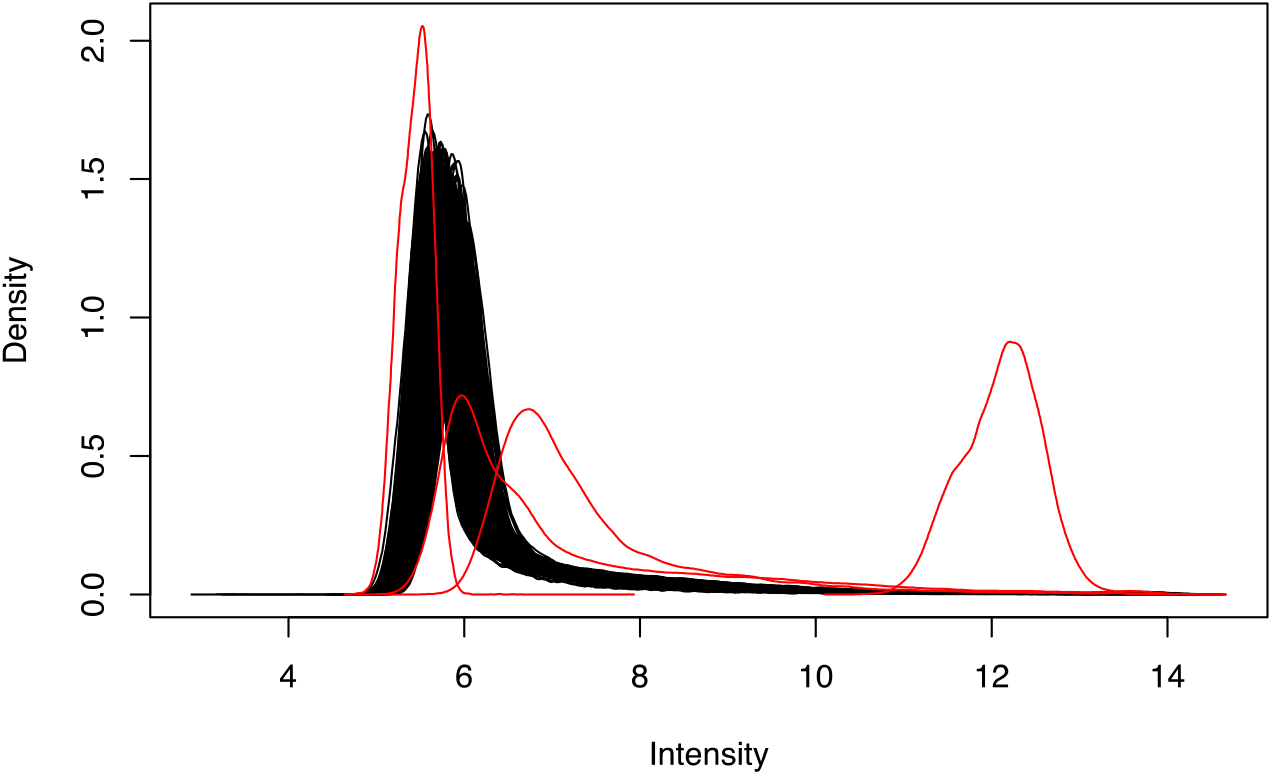
*Densities of gene expression intensity across the arrays of the four SOP outliers (in red) along with the densities of the rest of the data (black lines).*

We remove the four observations we consider technical outliers and compare with a fully-automated approach where we remove all suggested outliers without looking at them. This is done in one round with a cutoff of two standard deviations for all three methods of boxplots, PCA, and MA-plots. Accepting all outliers results in the removal of 59 observations. We also compare against removing no outliers. We refer to these three approaches as manual-, automatic-, and no outlier removal below.

**Fig. 8** below shows our results. There are 18 experiments in total. There are three outlier removal strategies: manual removal, no removal, and automatic removal. There are three dataset sizes: all data, half of the data, and 10% of the data. Finally, there are two hypothesis testing situations: one where we expect no difference between groups (denoted as the permutation situation), and one where we do expect differences (the bootstrap situation) for some genes. We show the usual type of boxplot: median, quartiles, and whiskers that extend to the outmost points no farther away from the median than 1.5 times the interquartile range.

**Fig. 8.**
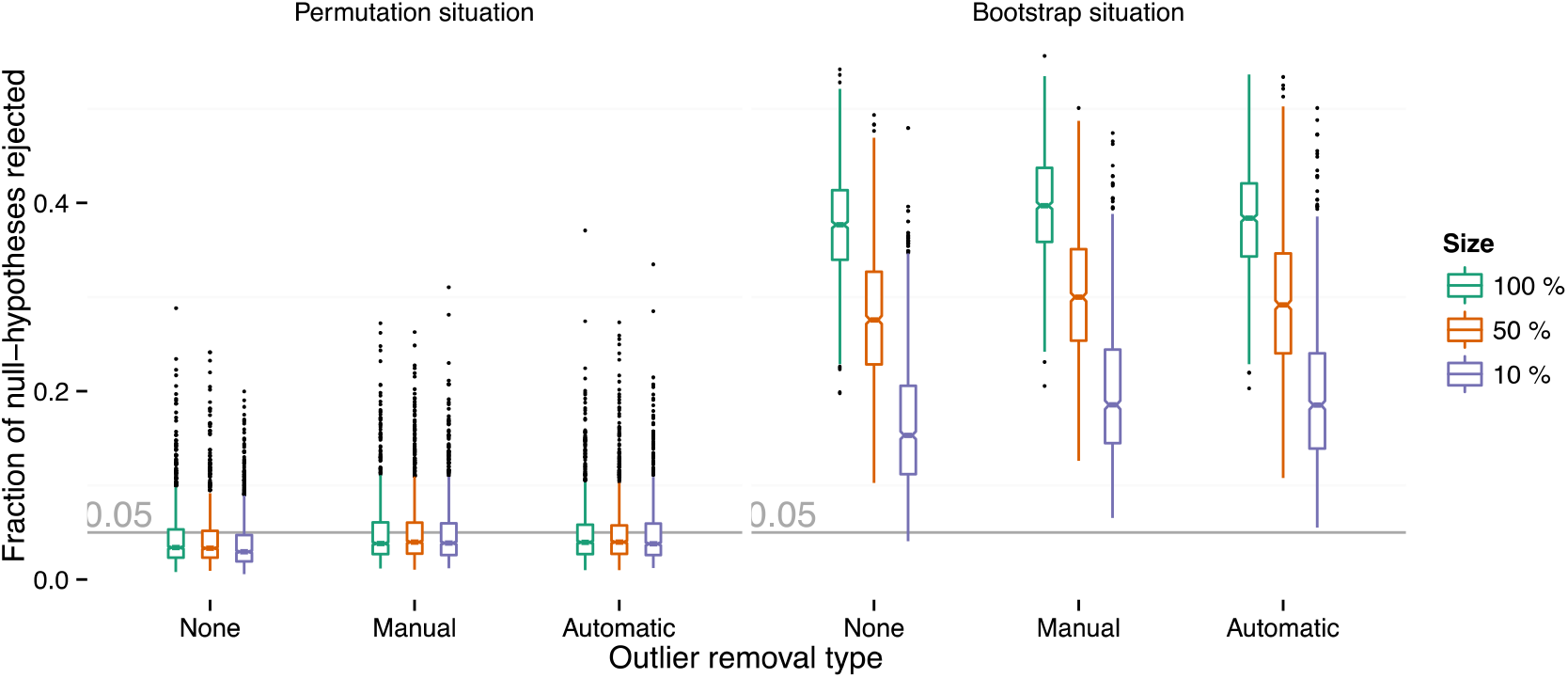
*Fraction of null-hypotheses rejected at a 5% significance level. The boxplots are the standard kind, with whiskers extending to the most extreme points within 1.5 times the interquartile range from the median. We have examined three different data sizes: all 832 observations of our data (green boxes), half of these data (orange), and 10% of them (violet). There are three different approaches to outlier removal: no outlier removal, manual removal, and automatic removal. We have examined rejection rates in two situations, one where the null hypothesis of no difference between genes is true, and one where we expect to observe a difference between the two groups for some genes. We see slightly improved error rate calibration and increased power for manual outlier removal in all cases.*

In the permutation situation there is no clear advantage to any method, though the no outlier removal strategy has a slightly lower *θ* than expected, which is especially clear for the smallest dataset. Both manual and automatic outlier removal push this value closer to the 0.05 error rate you would expect. We suspect we still get a slightly lower error rate than 0.05 due to dependence between genes.

In the bootstrap situation careful, manual outlier removal improves power over no outlier removal for all dataset sizes. This effect goes away for the automatic removal as you start removing useful information. For the smallest dataset, any removal is better than none at all.

All in all, there is some evidence that manual outlier removal increases power and that it calibrates your error rate under the null.

## 4 Discussion

### 4.1 Conclusions

This paper describes the NOWAC standard operating procedure for the removal of technical outliers. We have described the methods we use and provide an R-package implementation at https://github.com/3inar/nowaclean. By defining a common set of methods and lab measure cutoffs to detect and evaluate technical outliers, we believe we ensure greater consistency in the preprocessing of large sample microarray data sets. Further, by providing a detailed stand-alone documentation of how we do this, we believe we make it easier to understand and reproduce the research conducted.

### 4.2 Future work

As future work, we plan to improve the R package. Specifically, at the time of writing, some of the functions need better documentation. We would also like to provide interactive reports in Shiny, to make the SOP faster and easier to apply. These changes will be committed to the *nowaclean* git repository. We will eventually submit the package to CRAN or Bioconductor.

The microarray as a platform seems to be on its way out, and it seems likely that there will be a general move toward using RNA-Seq instead. We are uncertain to what extent our approach is applicable in an RNA-Seq setting. The lab measures will certainly change, and sequencing data is in its purest form count data. We need to provide an evaluation of this in the near future.

Finally, we plan to investigate whether outlier removal should be performed on a dataset-by-dataset basis, as we currently do it, of whether it would be better to merge all our datasets and do the outlier removal for this large combined dataset. The latter may have a smaller chance of removing valuable biological outliers.

## 5 Supporting Information

Online Resource 1: Full report of outlier removal in the demo data set.

Online Resource 2: The source code for the package version used in Online Resource 1.

We have made the demo dataset and our experiment code available online as http://dx.doi.org/10.18710/FGVLKS.

## 6 Acknowledgements

We are thankful to and impressed by the women that donated blood for this cancer research project. Bente Augdal, Merete Albertsen, and Knut Hansen were responsible for all infrastructure and administrative issues. Kajsa Møllersen has provided many useful comments to the SOP.

## 7 Conflict of interest

### Funding

This study was supported by a grant from the European Research Council (ERC-AdG 232997 TICE). The funders had no role in the design of the study; in the collection, analyses and interpretation of the data; in the writing of the manuscript; or in the decision to submit for publication.

Some of the data in this article are from the Cancer Registry of Norway. The Cancer Registry of Norway is not responsible for the analysis or interpretation of the data presented.

Microarray service was provided by the Genomics Core Facility, Norwegian University of Science and technology, and NMC – a national technology platform supported by the functional genomics program (FUGE) of the Research Council of Norway.

## 8 Statement of human rights

### Ethical approval

For this type of study formal consent is not required.

The NOWAC study was approved by the Norwegian Data Inspectorate and the Regional Ethical Committee of North Norway (REK). The linkages of the NOWAC database to national registries such as the Cancer Registry of Norway and registries on death and emigration from Statistics Norway was approved by the Directorate of Health. The women were informed about these linkages. Furthermore, the collection and storing of human biological material was approved by the REK in accordance with the Norwegian Biobank Act. Women were informed in the letter of introduction that the blood samples would be used for gene expression analyses.

